# Functional Contribution of Multienzyme Glucosome Condensates to Cellular Redox Homeostasis in Cancer Cells

**DOI:** 10.64898/2026.09.02.748920

**Authors:** Ashesh Sharma, B. K. Bijaya, Minjoung Kyoung, Songon An

## Abstract

Glucosomes are liquid-liquid phase-separated condensates observed in human cells, formed by phosphofructokinase and other rate-determining enzymes in glycolysis and gluconeogenesis. While glucosomes are spatially formed into small-, medium-, and large-sized assemblies in cancer cells, medium-sized glucosomes are functionally characterized to shunt glucose flux to the pentose phosphate pathway (PPP). As the PPP is the primary pathway responsible for maintaining cytosolic NADPH levels during oxidative stress, we hypothesize that medium-sized glucosomes regulate cellular redox homeostasis through the promotion of the PPP. In this work, we started treating Hs578T cells with hydrogen peroxide (H_2_O_2_) to evaluate how glucosomes respond to redox perturbation. High-content imaging demonstrated that H_2_O_2_ significantly promotes medium-sized glucosomes at both single-cell and population levels. The extracellular acidification rate by Seahorse extracellular flux analysis then corroborated that H_2_O_2_ effectively diverts glycolytic flux to the PPP through the upregulation of medium-sized glucosomes. We then investigated the glutathione redox cycle as a potential mechanistic link between medium-sized glucosomes and H_2_O_2_ detoxification. Treatment with oxidized glutathione (GSSG), but not reduced glutathione (GSH), markedly increased the population of cells showing medium-sized glucosomes. Moreover, shRNA-mediated knockdown of glutathione reductase, which converts GSSG to GSH at the expense of NADPH, attenuated H_2_O_2_-induced glucosome formation in Hs578T cells. Collectively, we demonstrate that glucosome-mediated metabolic reprogramming couples glucose metabolism to the glutathione redox cycle to facilitate H₂O₂ detoxification, thereby establishing the functional role of glucosomes in cellular redox homeostasis.

## Introduction

Cellular metabolism is orchestrated as a network of a large number of pathways for the breakdown (i.e., catabolism) and synthesis (i.e., anabolism) of biomolecules to maintain cell viability [1]. Among them, glucose metabolism is a centrally located metabolic network that primarily includes glycolysis, gluconeogenesis, the pentose phosphate pathway (PPP), and serine biosynthesis [1–3]. Glycolysis and its reciprocal pathway, gluconeogenesis, catalyze the interconversion between glucose and pyruvate. Meanwhile, the PPP uses glucose-6-phosphate to synthesize ribulose-5-phosphate while serine biosynthesis uses 3-phosphoglycerate to synthesize serine. Accordingly, their relative activities would depend in part on substrate availability, which is largely controlled by the rate-determining enzymes of the pathways. Ultimately, the regulation of these rate-determining enzymes allows a cell to balance its catabolic demands with its anabolic needs using glucose as both the source of energy and carbon [4–6].

Particularly, the PPP is an essential metabolic pathway that supports cellular reduction/oxidation (redox) homeostasis and/or nucleic acid biosynthesis, depending on metabolic demands [7]. Briefly, the PPP is divided into two branches [8]. The oxidative branch catalyzes the one-way synthesis of ribulose-5-phosphate from glucose-6-phosphate [7–9] whereas the non-oxidative branch catalyzes the interconversion among various sugars [8,10], including ribose-5-phosphate for nucleotide metabolism, and fructose-6-phosphate and glyceraldehyde-3-phosphate for glycolysis and gluconeogenesis [8]. Importantly, the oxidative PPP is a major source of cytosolic NADPH [8,11–13], accounting for 30-50% of NADPH in proliferating cells [14]. Even the non-oxidative branch allows for shuttling fructose-6-phosphate and glyceraldehyde-3-phosphate back into gluconeogenesis and subsequently the oxidative PPP for the maximal regeneration of NADPH [8,15]. In short, the PPP plays a critical role in maintaining NADPH levels in cells and allows for greater flexibility in cellular redox response under oxidative stress.

The pool of NADPH provides a cell with reducing power to maintain cellular redox homeostasis [11]. Cellular metabolism has been known to produce various reactive oxygen species (ROS), such as hydrogen peroxide (H_2_O_2_), superoxide, hydroxyl radical, peroxyl radical, and alkoxyl radical [16,17]. Among them, H_2_O_2_ is particularly notable as a persistent cellular ROS due to its relative stability, membrane permeability, and its role as a key mediator of redox signaling [16,17]. When a cell is exposed to >20 µM concentrations of H_2_O_2_, the glutathione redox cycle and catalase become two predominant regulators against H_2_O_2_ [18,19]. The glutathione redox cycle consists of two key enzymes: glutathione peroxidase (GPX) and glutathione reductase (GSR). GPX enzymatically reduces H_2_O_2_ to water while oxidizing reduced glutathione (GSH) to oxidized glutathione (GSSG). GSR then reduces GSSG back to GSH through the oxidation of NADPH [20]. Catalase (CAT), on the other hand, reduces H_2_O_2_ using heme as a cofactor [21]. However, mammalian CAT, including human CAT (hCAT), is sensitive to H_2_O_2_ and thus prone to be inactivated during its catalytic turnover [21–23]. To prevent such inactivation, human enzyme is known to tightly bind NADPH to maintain its catalytic activity [21,23,24]. Hence, cytosolic NADPH, primarily generated by the PPP, plays essential roles as both a redox cofactor and a protective molecule, emphasizing the significance of studying potential mechanisms of how glucose metabolism is metabolically regulated between glycolysis and the PPP in cells.

Meanwhile, various metabolic enzymes in glucose metabolism have been demonstrated to form liquid-liquid phase-separated condensates, namely the glucosomes, in human cells [25–28]. Glucosomes consist of at least four rate-determining enzymes in glycolysis and gluconeogenesis: phosphofructokinase liver-type (PFKL), fructose-1,6-bisphosphatase liver-type, pyruvate kinase muscle isoform 2 (PKM2), and phosphoenolpyruvate carboxykinase 1 [25]. At a single-cell level, glucosomes are spatially observed to form various sizes and are categorized into three subgroups: small- (< 0.1 µm^2^), medium- (0.1-3 µm^2^), and large-sized (3-8 µm^2^) glucosomes [25]. Among them, the formation of medium-sized glucosomes is significantly promoted at both single-cell and population levels in the presence of fructose-1,6-bisphosphate [25], excess of which is known to inhibit the bottle-neck step of glycolysis and thus shunt glycolytic flux into the PPP [29–33]. Importantly, methylene blue, which is well known to deplete the cellular NADPH level and thus upregulates the PPP [7,34–38], is also demonstrated to upregulate the formation of medium-sized glucosomes in HeLa and Hs578T cells [25]. Given the metabolic function of the PPP under oxidative stress, we have hypothesized the functional association of medium-sized glucosomes with NADPH-dependent processes such as cellular redox homeostasis.

Here, we have investigated how medium-sized glucosomes regulate cellular redox homeostasis through the promotion of PPP. We first employed fluorescence live-cell imaging strategies to quantitatively monitor the upregulation of medium-sized glucosomes in response to exogenous H_2_O_2_ treatment in various conditions. Seahorse metabolic flux measurements then provided additional evidence that H_2_O_2_ treatment redirects glucose flux from glycolysis toward the PPP. Importantly, when cellular redox balance was perturbed with exogenous glutathione (i.e., GSH and GSSG, respectively), medium-sized glucosomes were regulated accordingly to maintain homeostasis, demonstrating a mechanistic coupling between the glutathione redox cycle and medium-sized glucosome. Subsequent shRNA knockdown approaches further validated the coordination of glucosomes with the glutathione redox cycle, but the functional participation of hCAT in H_2_O_2_-induced glucosome formation was negligible. Collectively, we have provided compelling evidence of the functional coupling between the glutathione redox cycle and medium-sized glucosomes under H_2_O_2_-induced oxidative stress, revealing the functional significance of glucosomes in cellular redox homeostasis.

## Results

We have previously shown that human cancer cells upregulate medium-sized glucosomes in response to methylene blue [25], a small molecule that perturbs cellular redox homeostasis through the disruption of the NADPH pool [7,34–38]. However, a mechanistic association of medium-sized glucosomes with cellular redox homeostasis has not yet been elucidated. Therefore, in this work, we have investigated a hypothesis that the assembly of medium-sized glucosomes is functionally and mechanistically coupled with redox homeostasis by facilitating glucose flux into the PPP.

### Upregulation of medium-sized glucosomes in response to hydrogen peroxide (H_2_O_2_)

To test this hypothesis, we used fluorescence live-cell imaging to quantitatively understand the real-time response of medium-sized glucosomes in relation to changes in redox homeostasis in human cancer cells. The glucosome marker PFKL, tagged with a monomeric enhanced green fluorescent protein (PFKL-mEGFP), was transfected into human Hs578T breast cancer cells. We then profiled glucosome formation based on the prevalence of a predefined glucosome size at a single-cell level as we have previously categorized into three different subgroups: small-, medium- , and large-sized glucosomes [25,39]. When Hs578T cells expressing PFKL-mEGFP were treated with 100 µM H_2_O_2_, it was apparent that H_2_O_2_ robustly upregulated the formation of medium-sized glucosomes within an hour at both single-cell and population levels (**Figure 1A-B and 1E**).

**Figure 1.**
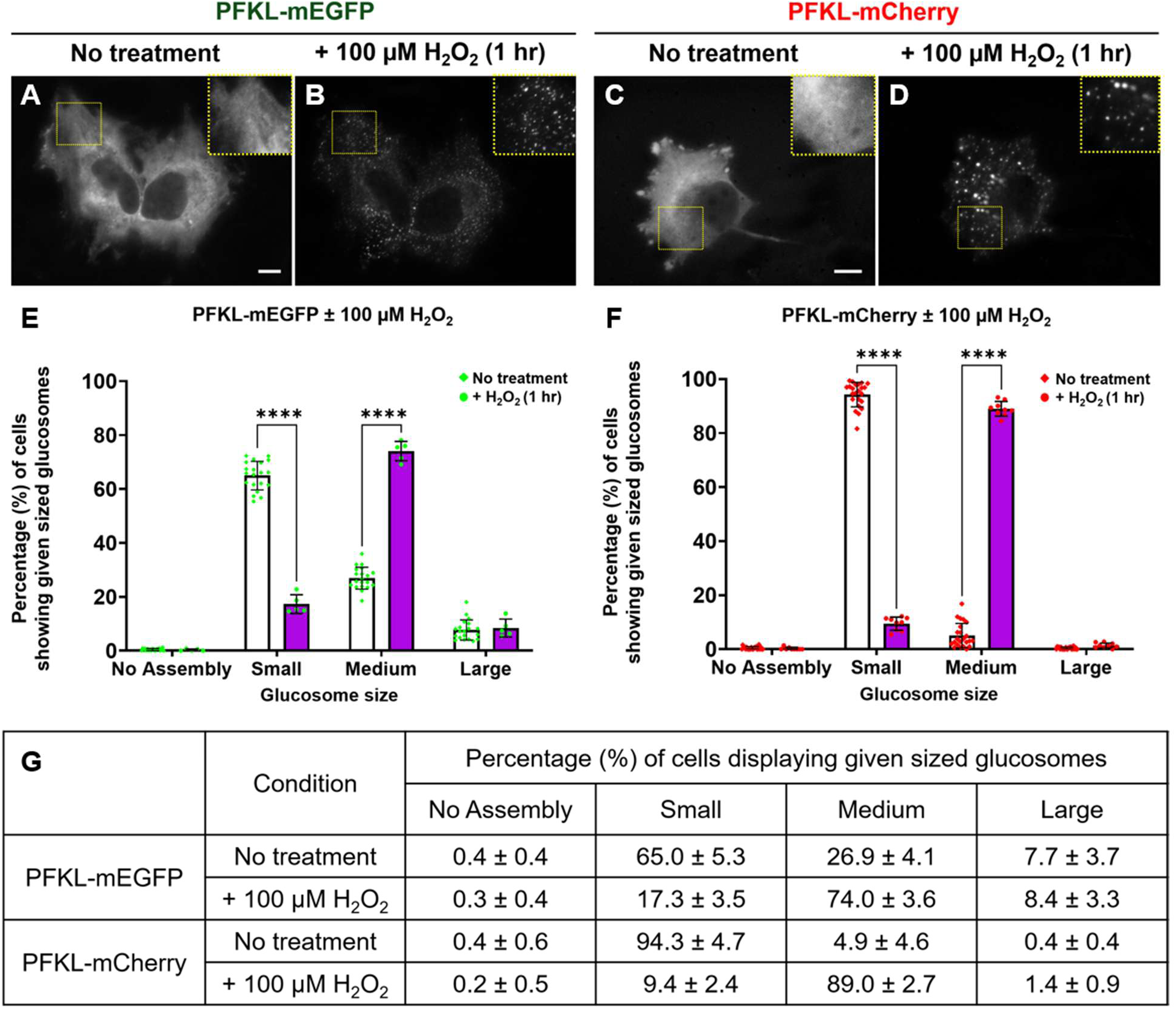
High-content imaging of glucosomes in response to H_2_O_2_. *A–D*, representative Hs578T cells expressing PFKL-mEGFP (A and B) and PFKL-mCherry (C and D) were imaged before (A and C) and after (B and D) the addition of H_2_O_2_ (100 µM, 1 hr). Insets indicate digitally magnified regions of interest. Scale bars, 10 µm. *E and F*, the percentages (%) of cells showing given sized glucosomes with PFKL-mEGFP and PFKL-mCherry, respectively, were quantified before (blank bars, *N_expt_* = 20 and *N_expt_* = 23, respectively) and after (colored bars, *N_expt_* = 5 and *N_expt_* = 9, respectively) the addition of H_2_O_2_ (100 µM, 1hr). Error bars represent standard deviations (SD). *G*, a table shows the average percentages (%) of Hs578T cells showing given sized glucosomes for each fusion protein, with or without the treatment of H_2_O_2_, along with their SDs (±). Statistical analyses were performed using Tukey’s multiple comparison tests for two-way ANOVA analysis. \*\*\*\**p <* 0.0001; ns, not significant; *N_expt_*, the number of independent experiments. H_2_O_2_, hydrogen peroxide; mEGFP, monomeric enhanced green fluorescent protein; mCherry, monomeric cherry fluorescent protein; PFKL, phosphofructokinase liver-type.

Alternatively, we also used another fusion protein of PFKL, tagged with a monomeric cherry fluorescent protein (PFKL-mCherry), to show that H_2_O_2_-induced formation of medium-sized glucosomes is not dependent on the fused fluorescent protein tag to PFKL (**Figure 1C-D and 1F**). Although mCherry itself, but not mEGFP, seems to possess an undetermined ability to dissolute biomolecular condensates in living cells [40], the mCherry fusion construct in our work did not suppress the H_2_O_2_-induced responses but rather promoted the formation of medium-sized glucosomes to a greater extent in Hs578T cells (i.e., ∼74% vs. ∼89%, **Figure 1E vs 1F**). A simultaneous decrease in the percentage of cells showing small-sized glucosomes was clearly monitored in both conditions (**Figure 1E-G**). In addition, we treated Hs578T cells expressing a mEGFP-tagged mutant PFKL, PFKL(N702T)-mEGFP, with 100 µM H_2_O_2_, which is catalytically active but unable to organize glucosomes in living cells [41]. In this case, however, we observed no discernible response after the treatment of 100 µM H_2_O_2_ (**Figure S1**). It appears that the H₂O₂-induced formation of medium-sized glucosomes represents a biologically relevant response of glucosomes to H₂O₂ treatment, suggesting a potential role of glucosomes in cellular redox homeostasis. Collectively, we demonstrate that Hs578T cells robustly respond to the treatment of H_2_O_2_ by upregulating the formation of medium-sized glucosomes.

### Metabolic shunt of glycolytic flux to the PPP in response to H_2_O_2_

Next, we measured real-time extracellular acidification rates (ECAR) using Seahorse Extracellular XF Flux Analyzer to determine whether H_2_O_2_-induced glucosome formation is associated with metabolic changes in glucose metabolism. Real-time ECAR measurements following an acute treatment of 100 µM H_2_O_2_, relative to vehicle (i.e., H_2_O), showed a gradual decrease in ECAR signals (**Figure 2**), indicating a real-time reduction of glycolytic flux. To evaluate whether the H_2_O_2_-induced reduction in ECAR is due to changes in mitochondrial respiration, we subsequently treated the cells with 0.5 µM mitochondrial respiration complex inhibitors (i.e., a mixture of rotenone and antimycin A [42,43]). Although they efficiently blocked oxygen consumption rates in mitochondria (**Figure S2**), no change in ECAR was observed (**Figure 2A**), suggesting that the H_2_O_2_-induced shunt of glucose flux appears to be mitochondria-independent under our conditions. When we further treated the cells with 50 mM 2-deoxyglucose (2-DG), we observed a sharp decrease in ECAR (**Figure 2A**), verifying that the ECAR effects that we had observed were due to the changes in glycolytic flux. Given metabolomics studies revealing H_2_O_2_-induced promotion of the PPP in mammalian cells [44–47], our ECAR kinetic profiling (**Figure 2**) is in good agreement with glucose being shunted from glycolysis to the PPP. Therefore, our results show that H_2_O_2_-treated Hs578T cells promoting medium-sized glucosomes effectively divert glycolytic flux to the PPP in response to the perturbation of cellular redox homeostasis.

**Figure 2.**
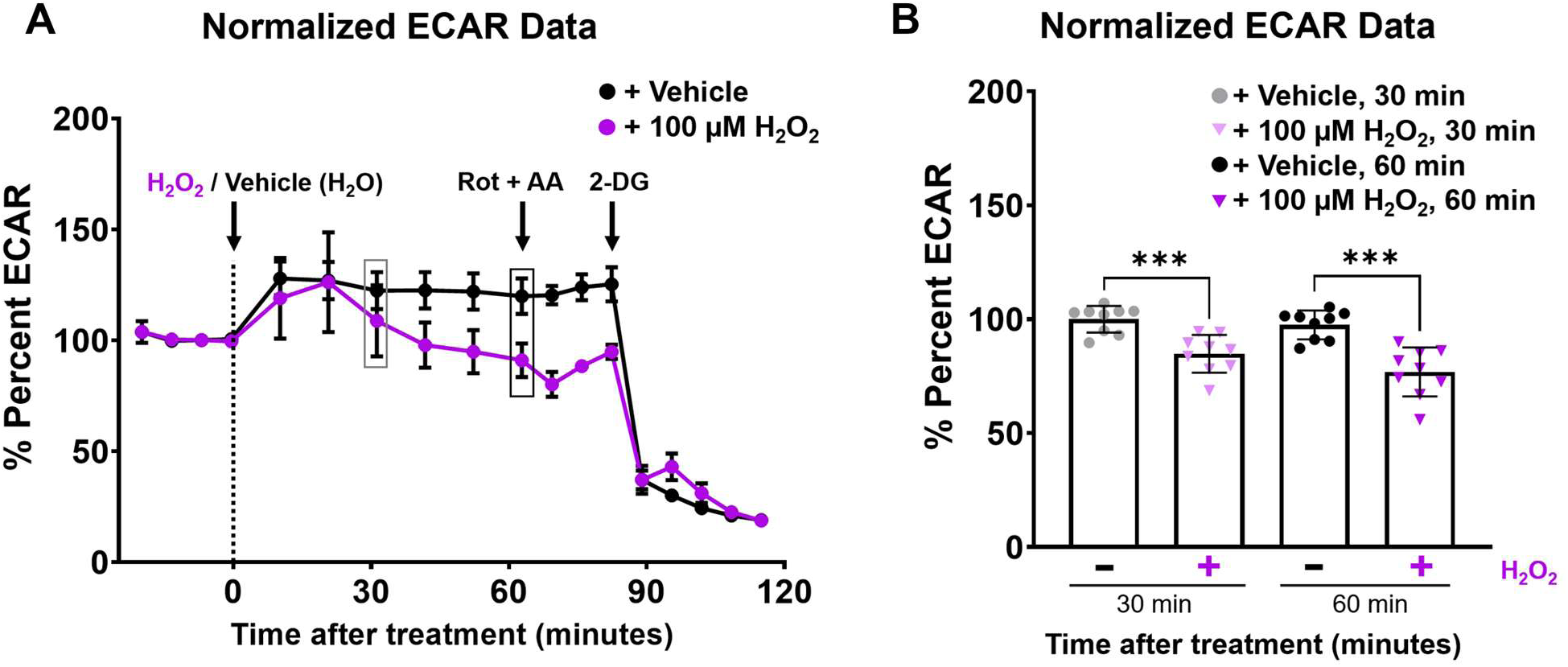
Seahorse extracellular flux analysis of Hs578T cells with H_2_O_2_. *A,* normalized percentage of extracellular acidification rate (% ECAR) from Hs578T cells treated with H_2_O_2_ (purple line) shows a gradual decrease after 20 min of H_2_O_2_ treatment relative to water-treated control cells (black line). Black arrows indicate the timepoint for the addition of i) 100 µM H_2_O_2_ or vehicle (H_2_O), ii) a mixture of 0.5 µM rotenone (Rot) and 0.5 µM antimycin A (AA), and iii) 50 mM 2-deoxyglucose (2-DG). Each data point represents the average % ECAR signals from three technical replicates of three independent trials, with error bars representing standard deviations (SD). *B,* extracted normalized % ECAR signals of H_2_O_2_-treated Hs578T cells (purple triangle) and water-treated control cells (black circle) at 30 and 60 min. A bar graph represents the average % ECAR signals from three technical replicates of three independent trials with SDs. Statistical analyses were performed using Student’s *t*-test with Welch’s correction. \*\*\**p* < 0.001.

### Functional association of the glutathione redox cycle with glucosomes

To investigate a mechanism behind the functional association of glucosome assemblies with H_2_O_2_-induced redox perturbation, we hypothesized that glucosome assemblies may be directly coupled to the glutathione redox cycle by shunting glycolytic flux to the PPP and thus increasing NADPH in a cell (**Figure 3A**). In this work, we first employed oxidized glutathione (GSSG) to perturb the glutathione homeostasis. Hs578T cells expressing PFKL-mEGFP were treated with 10 mM GSSG or vehicle (0.3 mM HEPES, pH 7.4). After an hour, subsequent glucosome profiling showed a significant (*p* < 0.0001) increase in the number of cells showing medium-sized glucosomes when compared with vehicle-treated cells (**Figure 3**). This result indicates that GSSG-induced perturbation appears to couple the reduction of GSSG with the oxidation of NADPH, thereby promoting the PPP through the formation of medium-sized glucosomes. Conversely, when Hs578T cells were treated with 10 mM reduced glutathione (GSH) for an hour, no change was detected. Even after 16 hr, only a small increase in the number of cells showing small-sized glucosomes (*p* < 0.0001) was seen, but no increase in the number of cells showing medium-sized glucosomes was monitored relative to vehicle-treated controls (**Figure 4**). This implies that the treatment of reduced GSH perturbing glutathione homeostasis is not directly associated with either the PPP or medium-sized glucosomes. Taking together, our results suggest that the glutathione redox cycle is mechanistically coupled to the PPP-associated NADPH production through the formation of medium-sized glucosomes.

**Figure 3.**
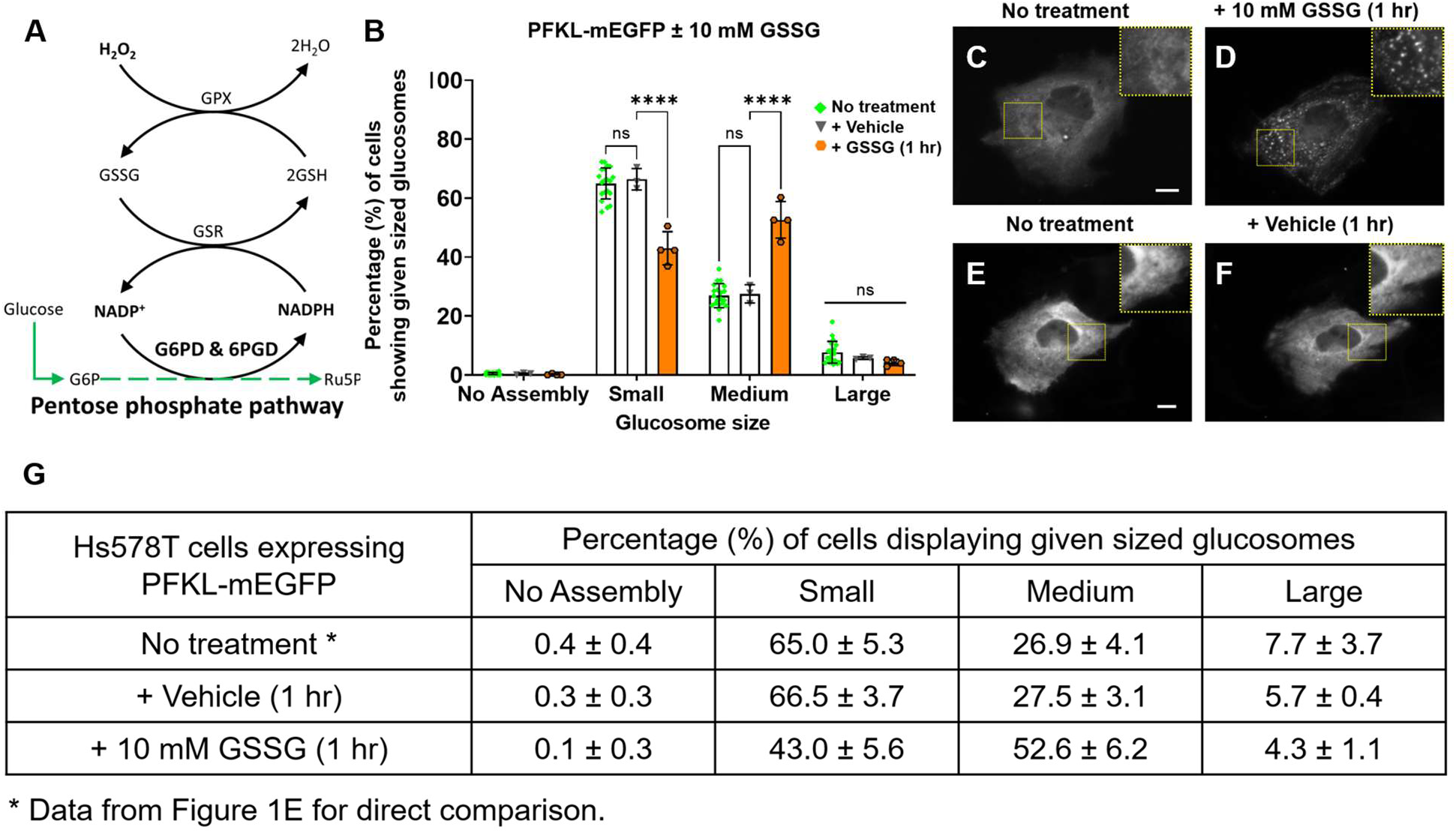
Effect of oxidized glutathione (GSSG) on glucosomes in Hs578T cells. *A*, a schematic diagram of the glutathione redox cycle shows detoxification of H_2_O_2_ by glutathione peroxidase (GPX) using reduced glutathione (GSH) and glutathione reductase (GSR) using NADPH, which is produced by glucose-6-phosphate dehydrogenase (G6PD) and 6-phosphogluconate dehydrogenase (6PGD) in the pentose phosphate pathway (PPP). G6P, glucose-6-phosphate; Ru5P, ribulose-5-phosphate. *B,* the percentages (%) of Hs578T cells showing given sized glucosomes were quantified before and after the addition of GSSG (10 mM) (*N_expt_* = 4) and vehicle (0.3 M HEPES) (*N_expt_* = 3). Note that the ‘No treatment’ data from Figure 1E is also included in this graph as ‘No treatment’ for direct comparison. Error bars represent standard deviations (SD). *C–F*, representative Hs578T cells expressing PFKL-mEGFP were imaged before (C and E) and after the treatment of 10 mM GSSG (D) and vehicle (F). Scale bars, 10 µm. Insets indicate digitally magnified regions of interest. *G*, a table shows the average percentages (%) of Hs578T cells showing given sized glucosomes with or without GSSG, along with their SDs (±). Statistical analyses were performed using Tukey’s multiple comparison tests for two-way ANOVA analysis. \*\*\*\**p* < 0.0001; ns, not significant; *N_expt_*, the number of independent experiments.

**Figure 4.**
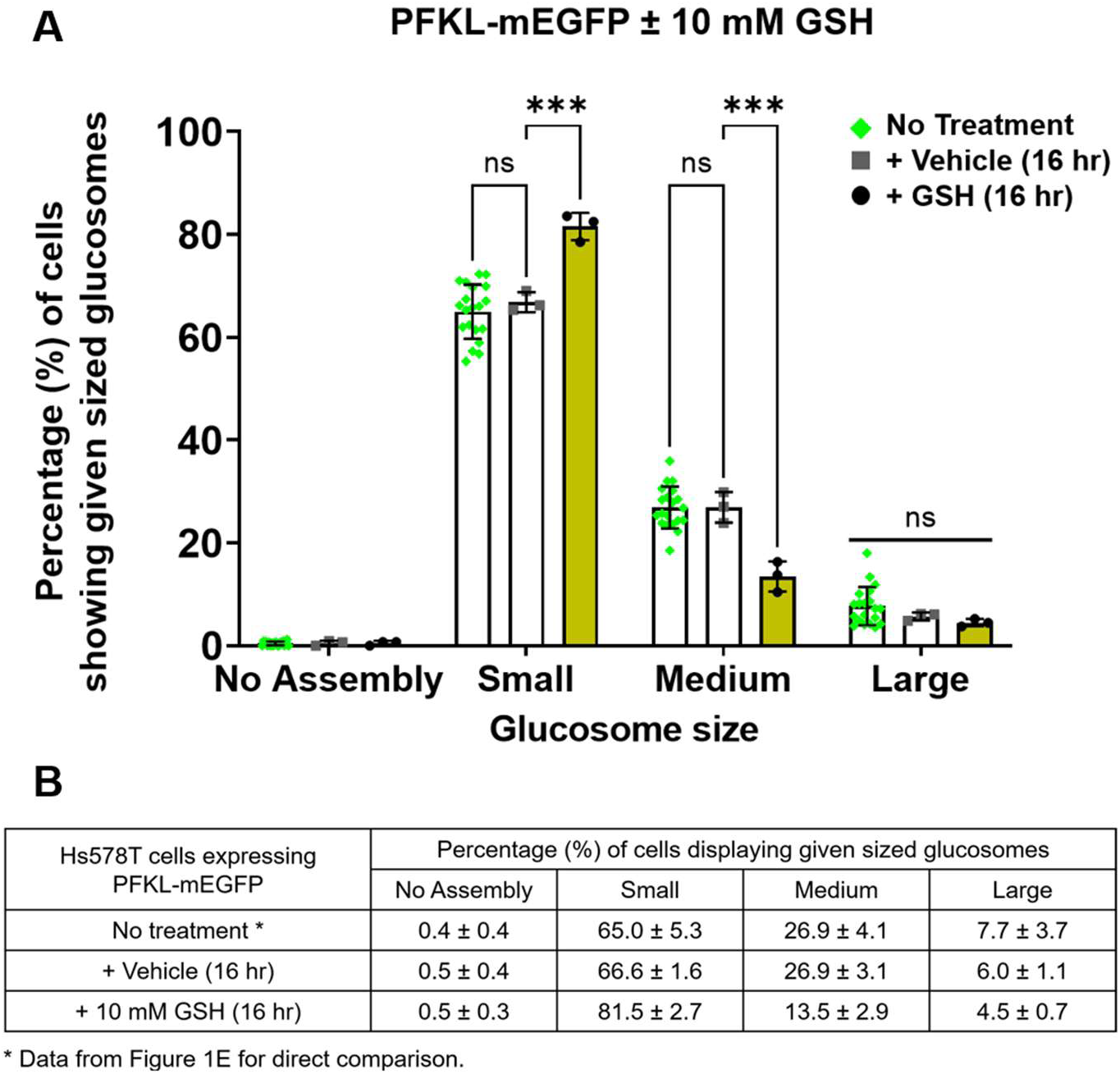
Effect of reduced glutathione (GSH) on glucosomes in Hs578T cells. *A,* the percentages (%) of Hs578T cells showing given sized glucosomes were quantified before and after the addition of GSH (10 mM, 16 hr) (*N_expt_* = 3) and vehicle (0.3 M HEPES) (*N_expt_* = 3). *B*, a table shows the average percentages (%) of Hs578T cells showing given sized glucosomes with or without GSH, along with error bars representing standard deviations (SD) (±). Note that the ‘No treatment’ data from Figure 1E is also included in this graph as ‘No treatment’ for direct comparison. Statistical analyses were performed using Tukey’s multiple comparison tests for two-way ANOVA analysis. \*\*\**p* < 0.001; ns, not significant; *N_expt_*, the number of independent experiments.

Next, we investigated glutathione reductase (GSR), which is responsible for operating the glutathione redox cycle by catalyzing the conversion of GSSG to GSH in human cells [48,49]. Since exogenous GSSG promoted the formation of medium-sized glucosomes (**Figure 3**), we hypothesized that human GSR (hGSR) would be mechanistically essential to coordinate the glutathione redox cycle with medium-sized glucosomes for cellular defense mechanisms against H_2_O_2_. We first conducted shRNA-mediated knockdown of hGSR in Hs578T cells expressing PFKL-mEGFP as a glucosome marker. The expression of shRNA_hGSR_ itself in Hs578T cells, indicated by the expression of an internal turboGFP marker, didn’t influence the formation of medium-sized glucosomes in Hs578T cells (**Figure 5A**). However, in the presence of 100 µM H_2_O_2_, we observed that the significantly smaller number of cells promoted medium-sized glucosomes when hGSR was knocked down, relative to controls expressing shRNA_Scramble_ or no shRNA (**Figure 5**). Given that western blot analysis showed ∼36 % knockdown of hGSR under our conditions (**Figure S3A-B**), this data strongly supports that hGSR is directly associated with the H_2_O_2_-induced formation of medium-sized glucosomes. Therefore, we demonstrate that the glutathione redox cycle is functionally coupled with NADPH-producing glucosomes through the action of GSR, which effectively detoxifies H_2_O_2_ (**Figure 3A**).

**Figure 5.**
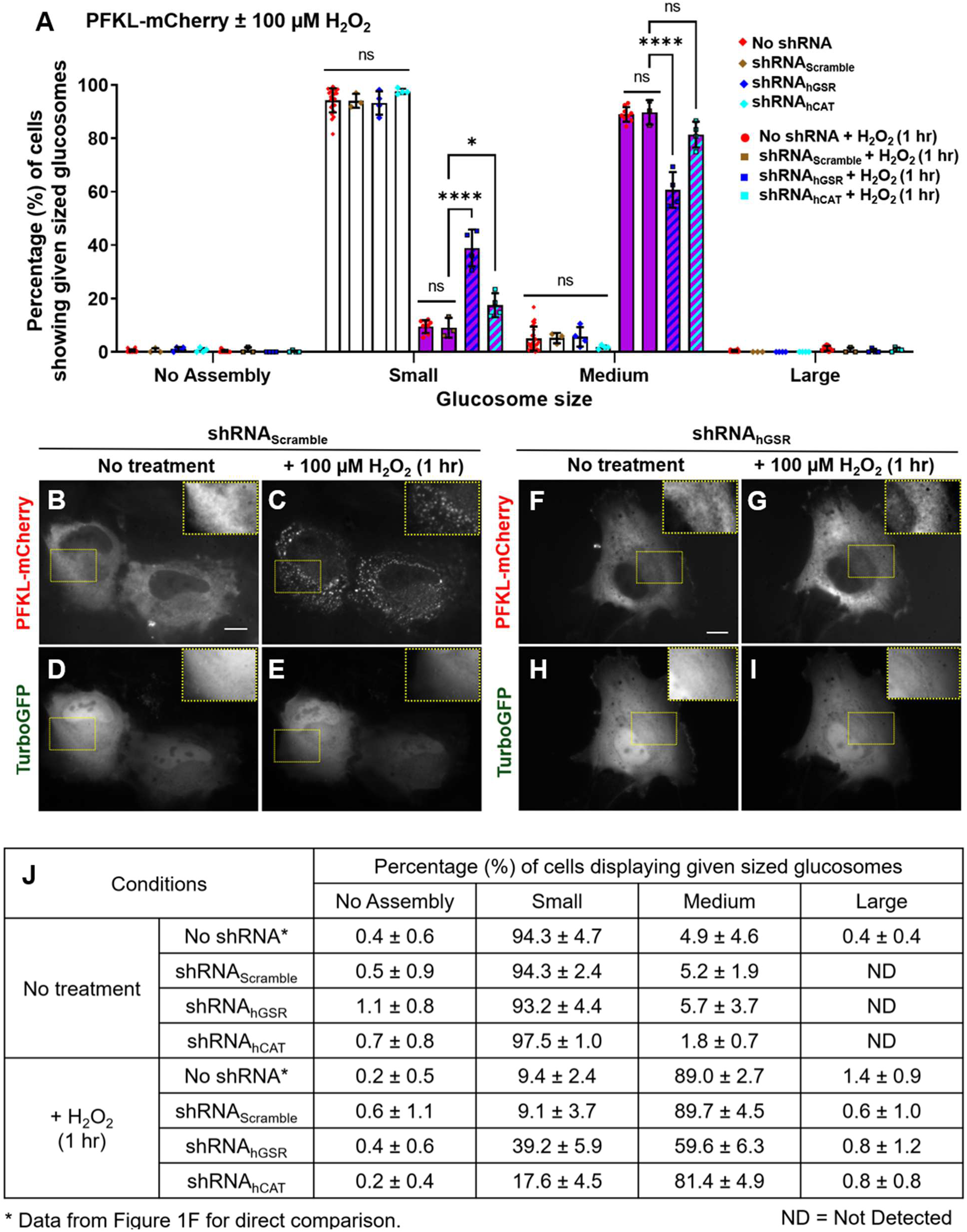
Knockdown effect of human glutathione reductase (hGSR) and catalase (hCAT) on H_2_O_2_-induced glucosomes in Hs578T cells. *A,* the percentages (%) of Hs578T cells showing given sized glucosomes were quantified before (shRNA_Scramble_, *N_expt_* = 3; shRNA_hGSR_, *N_expt_* = 4; shRNA_hCAT_, *N_expt_* = 4) and after (shRNA_Scramble_, *N_expt_* = 3; shRNA_hGSR_, *N_expt_* = 3; shRNA_hCAT_, *N_expt_* = 4) the addition of H_2_O_2_ (100 µM, 1 hr). Note that the data from Figure 1F are also included in this graph as ‘No shRNA’ and ‘No shRNA+ H_2_O_2_ (1 hr)’ for direct comparison. Error bars represent standard deviations (SD). *B-I,* representative Hs578T cells expressing PFKL-mCherry (B, C, F, and G) with shRNA_Scramble_ (B and C) and shRNA_hGSR_ (F and G) were imaged before (B and F) and after (C and G) the addition of H_2_O_2_ (100 µM, 1 hr). Insets indicate digitally magnified regions of interest. Expression of shRNA is confirmed by the expression of an internal marker, turboGFP, of the plasmid (D, E, H, and I). *J*, a table shows the average percentages (%) of Hs578T cells showing given sized glucosomes in various conditions tested here, along with their SDs (±). Statistical analyses were performed using Tukey’s multiple comparison tests for two-way ANOVA analysis. * *p* < 0.05, **** *p* < 0.0001; ns, not significant; *N_expt_*, the number of independent experiments.

### No participation of the catalase redox cycle in H_2_O_2_-induced glucosome formation

We then evaluated if human catalase (hCAT), which detoxifies H_2_O_2_ [21] and also consumes NADPH to remain catalytically active [23], has any contribution to the promotion of H_2_O_2_-induced medium-sized glucosomes in Hs578T cells. When we knockdown hCAT through the expression of shRNA_hCAT_ (**Figure S3C-D**), there was only a non-significant decrease in the number of cells showing H_2_O_2_-induced medium-sized glucosomes relative to controls (**Figure 5A and 5J**). To determine whether this decrease was due to reduced H_2_O_2_ levels resulting from the detoxifying activity of hCAT, we heterologously and stably expressed *E. coli* catalase (ecCAT) containing an *N*-terminal mEGFP tag in Hs578T cells, generating Hs578T-ecCAT cells. We then confirmed that heterologously expressed mEGFP-ecCAT in Hs578T cells appears to be functionally active because it effectively decreased the effect of 100 µM H_2_O_2_ by ∼57 % on the formation of medium-sized glucosomes (**Figure S4**). However, when we knockdown hCAT with shRNA_hCAT_ in the Hs578T-ecCAT cells, H_2_O_2_ did not trigger any noticeable change in the formation of medium-sized glucosomes (**Figure S5**), relative to controls, confirming no functional contribution of hCAT in H_2_O_2_-induced glucosome formation. Considering Hs578T-ecCAT cells expressing shRNA_hGSR_ showed a significant decrease in response to H_2_O_2_ treatment (**Figure S5**), this data reassures the major role of hGSR, but not hCAT, in the H_2_O_2_-induced formation of glucosomes.

## Discussions

We demonstrate that multienzyme glucosome assemblies coordinate with the glutathione redox cycle to promote cellular adaptation of glucose metabolism to oxidative stress in human cancer cells. Briefly, we show that cellular redox perturbations induced by the treatment of H₂O₂ (**Figure 1**) and methylene blue [25] promote the formation of medium-sized glucosomes, which are associated with the shunting of glycolytic flux toward the PPP (**Figure 2** and [25,50]). Direct perturbation of the glutathione redox cycle with and without H_2_O_2_ (**Figures 3, 5, and S5**) are also sufficient to modulate glucosome assemblies, strengthening their direct association. Therefore, this study identifies medium-sized glucosomes as functionally active metabolic hubs that coordinate cellular metabolic response to oxidative stress.

In human cells, enzymatic detoxification of H_2_O_2_ is mediated by several enzymes, including peroxiredoxin (PRX), catalase (CAT) and glutathione peroxidase (GPX), in a concentration-dependent manner. When the intracellular concentration of H_2_O_2_ rises, an ubiquitous class of thiol-dependent two-cysteine peroxidases (i.e., PRX) is turned on to maintain H₂O₂ concentrations at the range of ∼1–10 nM [16,19,51]. However, when cells are exposed to 20-100 µM H_2_O_2_ [52–54], the active cysteines of PRX are hyperoxidized and thus inactivated [18,53]. Consequently, at such high micromolar ranges, CAT and GPX are proposed to play key roles in regulating intracellular H₂O₂ levels [18,19]. CAT, a heme-containing peroxisomal enzyme that detoxifies H_2_O_2_ [21], is localized primarily in peroxisomes. Under oxidative stress, however, it is retained in the cytoplasm [55] and found to bind NADPH to protect itself against deactivation [23]. In parallel, GPXs are involved in H_2_O_2_ detoxification, along with GSR, in the glutathione redox cycle, which requires NADPH as a reducing cofactor. Specifically, in human cells, there are eight isoforms of GPXs, which differ in their substrate selectivity [56]. For example, GPX1 and GPX2 exhibit similar selectivity for H_2_O_2_ and soluble hydroperoxide, whereas GPX4 shows substantial activity toward phospholipid hydroperoxides relative to H_2_O_2_ and soluble hydroperoxides [57]. Therefore, the active participation of the glutathione redox cycle in response to 100 µM H_2_O_2_ under our conditions appears pertinent to the cellular response, which we further elaborate here by revealing the functional contribution of glucosome assemblies as part of the cellular adaptation to oxidative stress.

Treatment of H_2_O_2_ has been also reported to promote the PPP flux in eukaryotic cells. In yeasts, untargeted metabolomics has shown that 2 mM H_2_O_2_ significantly increases about 2-fold the concentration of ribose-5-phosphate, the product of the oxidative PPP, relative to untreated cells [58]. Similarly, a ∼4.5-fold increase of ribose-5-phosphate is also observed from mouse embryonic fibroblasts with 150 µM H_2_O_2_ [44]. An untargeted metabolomics study in HepG2 cells treated with 5 mM H_2_O_2_ has also showed the increased levels of various PPP metabolites, including 6-phosphogluconolactone and ribulose-5-phosphate, over the course of 60 min of the treatment [46]. Furthermore, in human skin fibroblasts, time-resolved mass spectrometry has revealed the rerouting of glucose flux to the PPP within seconds of 500 µM H_2_O_2_ treatment [45]. Additional ^13^C labeling experiments, in the same study, have confirmed that H_2_O_2_ treatment promotes multiple recycling of the PPP metabolites through both oxidative and non-oxidative PPP pathways to maximize NADPH regeneration [45]. Meanwhile, a study using human HAP1 cells has also shown an increase in PPP-derived metabolites by the treatment of 100 µM H_2_O_2_ [47]. Taking all together with Seahorse metabolic flux data presented in this work (**Figure 2**) and the literature [47], our results strongly support that medium-sized glucosomes induced by H_2_O_2_ effectively increase the PPP flux to contribute to cellular redox homeostasis.

Meanwhile, glycolytic enzymes are also known to serve as redox-sensitive nodes, providing carbon and energy for the synthesis and regeneration of cellular antioxidants [59]. For instance, under high oxidative stress, like radiation exposure, human lymphoblastoid cells are shown to upregulate a tumor suppressor protein p53, which in turn upregulates an enzyme that inhibits PFK by depleting its allosteric activator, fructose-2,6-bisphosphate [60,61]. Meanwhile, PKM2 in human A549 lung cancer cells and glyceraldehyde-3-phosphate dehydrogenase in human HAP1 cells are found to be inactivated by ROS through the oxidation of functionally important cysteine residues [47,62,63]. Consequently, the inactivated glycolytic enzymes appear to direct glucose flux to the PPP [58,59], thereby counteracting increased cellular ROS and enhancing cancer cell survival [62]. Considering PFK and PKM2 are two of the known components of the glucosome in human cells [25], we propose that the multienzyme glucosome assemblies that regulate glucose metabolism [25,28,50,64–66] function as dynamic subcellular nodes that reprogram glycolytic flux to combat oxidative stress.

In conclusion, we propose a model (**Figure 6**), which describes a subcellular regulation mechanism of glucose flux via differently sized glucosomes in response to changes in redox homeostasis. Under basal conditions, small-sized glucosomes predominantly operate glycolysis near mitochondria [28], supporting energy generation and lactate synthesis [6,67–69]. During acute oxidative stress, however, the glutathione redox system primarily detoxifies H_2_O_2_, leading to the consumption of NADPH involving the activity of GSR. In response, glucosomes organize into medium-sized assemblies and shunt glucose flux to the PPP for the regeneration of NADPH. Collectively, we propose that medium-sized glucosomes function as dynamic metabolic entities that orchestrate the diversion of glycolytic flux to the PPP under oxidative stress, revealing the functional significance of glucosomes in cellular redox homeostasis.

**Figure 6.**
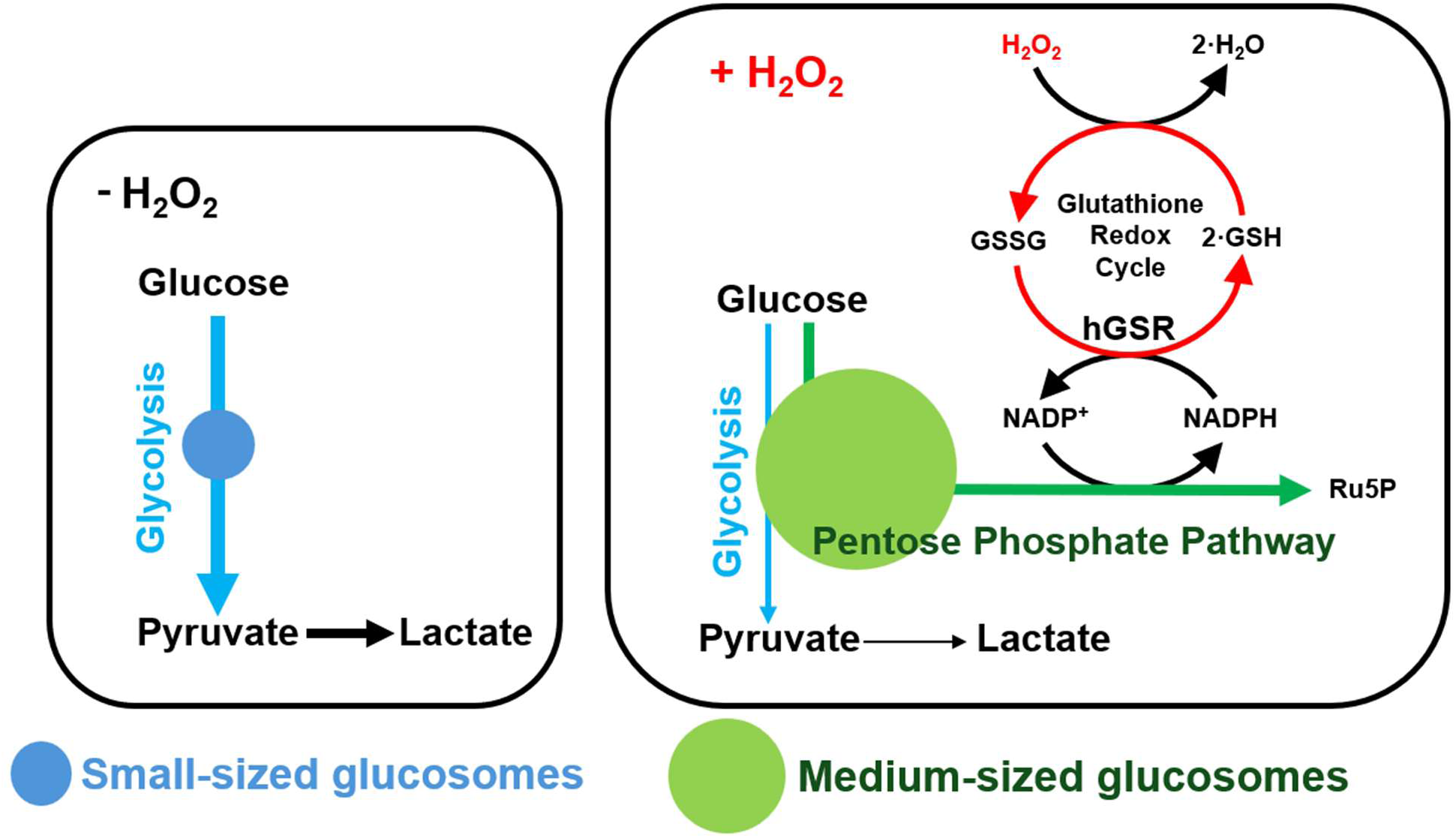
A schematic diagram of a proposed functional contribution of glucosomes to cellular redox homeostasis. Briefly, under basal conditions (i.e., in the absence of H_2_O_2_), small-sized glucosomes direct glucose to pyruvate and lactate (blue and black arrows) through glycolysis. Upon oxidative stress (i.e., in the presence of H_2_O_2_), the glutathione redox cycle (red arrows) detoxifies H_2_O_2_ by consuming NADPH. To support the detoxification process, medium-sized assemblies are formed and thus redirect glucose flux into the pentose phosphate pathway (green arrow) to regenerate NADPH.

## Materials and Methods

### Cloning and plasmid preparation

A plasmid expressing mEGFP-ecCAT was constructed in this work. Briefly, the gene expressing *E. coli* catalase (ecCAT) was amplified from pBad-HPII plasmid (Addgene, Cat. #105839) by PCR with a set of primers containing additional overhangs corresponding to XhoI and KpnI restriction sites. The ecCAT amplicon and the mEGFP-C1 vector (Clontech) were digested, respectively, with XhoI-HF (New England Biolabs (NEB), Cat. #R0146S) and KpnI-HF (NEB, Cat. #R3142S) and purified with Qiagen PCR purification kit (Qiagen, Cat. #28104) according to the manufacturer’s protocol. The digested and purified amplicon and vector were then subjected to ligation using T4 DNA Ligase (NEB, Cat. #M0202S), creating a plasmid expressing mEGFP-ecCAT under the CMV promoter. The ligated product was then transformed and amplified in chemically competent XL1-blue cells and purified using Thermo Scientific GeneJet plasmid miniprep kit (Thermo Scientific, Cat. #K0503). The resulting plasmid was verified by Sanger sequencing. Note that the plasmids expressing PFKL-mEGFP and PFKL-mCherry were available from previous works [25].

### Preparation of shRNA plasmids

Plasmids expressing short-hairpin RNA (shRNA) were constructed by ligating validated oligonucleotide sequences into the pGFP-C-ShLenti vector (Origene, Cat. #TR30023) using BamHI and BsmBI restriction sites. Briefly, a set of 5’-phosphate oligonucleotide sequences for human catalase (hCAT) [70] and glutathione reductase (hGSR) [71], respectively, were obtained from GenScript:

hCAT: 5’-GATCGTGCGGAGATTCAACACTGCCAATGATGATCAAGAGTCATCATTGGCAGTGTTGAATCTCCGCACTTTTTG-3’ and 5’-AGCTCAAAAAGTGCGGAGATTCAACACTGCCAATGATGACTCTTGATCATCATTGGCAGTGTTGAATCTCCGCAC-3’

hGSR: 5’-GATCATGTGAGCCGCCTGAATGCCATCTATCAATCAAGAGTTGATAGATGGCATTCAGGCGGCTCACATTTTTTG-3’ and 5’-AGCTCAAAAAATGTGAGCCGCCTGAATGCCATCTATCAACTCTTGATTGATAGATGGCATTCAGGCGGCTCACAT-3’

Each set of oligonucleotides were then subjected to annealing in annealing buffer (500 mM NaCl) using a custom PCR profile that allowed for a 1 °C/min cooling rate from 80 °C to 25 °C. Meanwhile, the vector was digested with BsmBI-v2 (NEB, Cat. #R0739S) and BamHI-HF (NEB, Cat. #R3136S) and purified using Qiagen PCR purification kit. The annealed oligos were then ligated with the double-digested pGFP-C-ShLenti vector using T4 DNA ligase, creating plasmids expressing shRNA_hCAT_ and shRNA_hGSR_, respectively. The ligated product was then transformed and amplified in chemically competent XL1-Blue cells and purified using Thermo Scientific GeneJet plasmid miniprep kit. The resulting plasmids were verified by DNA sequencing (plasmidsaurus). Note that a negative control plasmid expressing scrambled shRNA (i.e., shRNA_Scramble_) was purchased from Origene (Origene Cat. #TR30021).

### Cell culture

Hs578T cells (ATCC, Cat. #HTB-126) were initially grown in Dulbecco’s Modified Eagle’s Medium (DMEM, Cytiva, Cat. #SH30022.01) supplemented with 10% FBS (Sigma, Cat. #F2442), 1 mM sodium pyruvate (Sigma, Cat. #P5280), and 50 µg/mL gentamicin (Corning, Cat. #61-098-RF). Cells were maintained in a ThermoFisher HeraCell CO₂ incubator (37 °C, 5% CO₂, and 95% humidity) with subculturing every 3-4 days. After ∼2 weeks, the cells were transitioned into the Roswell Park Memorial Institute 1640 (RPMI 1640, Corning, Cat. #10-040-CV) medium containing 10% dialyzed FBS (dFBS), and 50 µg/mL gentamicin. Note that FBS was dialyzed against 0.9% NaCl at 4 °C for ∼48 hr with 12-14 KDa MWCO dialysis membrane (Spectrum, Cat. #S432706). After two weeks of adaptation to the RPMI 1640 growth medium with subculturing every 3-4 days, cells were deemed ready for experiments.

### Transfection

For live-cell imaging, Hs578T cells were harvested using a trypsin-EDTA solution (Corning, Cat. #25-053-Cl), and ∼125,000 cells were seeded into 35 mm glass-bottom petri dishes (MatTEK, Cat. #P35G-1.5-14-C) in the RPMI 1640 growth medium without gentamicin. ∼24 hr after seeding, cells were transfected with a plasmid coding for the indicated enzyme as previously described [25,39]. Briefly, 800-1600 ng of a plasmid and 1.5-3.0 µL of Lipofectamine 2000 (Thermo Scientific, Cat. #11668027) were diluted respectively in 50 µL of Opti-MEM™ I (Gibco, Cat. #11058) and incubated for 5 min at room temperature. Subsequently, the two solutions were gently mixed and incubated for additional 30 min. The transfection mixture was then gently diluted with 900 µL of Opti-MEM™ I and transferred to the MatTEK dish with cells in 1 mL of Opti-MEM™ I. The cells were placed in a CO_2_ incubator at 37 °C for ∼5-6 hr, after which they were incubated with the fresh, gentamicin-free RPMI 1640 growth medium at 37 °C in a CO_2_ incubator for ∼18-26 hr prior to imaging.

### Fluorescent live-cell imaging

Transfected cells were imaged at room temperature using Nikon Eclipse Ti inverted microscope using a 60x objective lens (Nikon CFI Plan Apo TIRF, 1.45 N.A.) as described before [25,39]. Briefly, cells were washed 3 times for 10 min each and then incubated for another 60 min at room temperature in buffered saline solution (BSS: 20 mM HEPES (pH 7.4), 135 mM NaCl, 5 mM KCl, 1.6 mM CaCl_2_, and 1 mM MgCl_2_). Cells were then visualized and categorized based on the presence of small-, medium-, or large-sized glucosomes as previously described [25]. Wide-field fluorescence imaging was performed using custom filter sets from Chroma Technology [25]. For mEGFP detection, a Z488/10-HC clean-up, HC TIRF dichroic, and 525/50-HC emission filter were used. For mCherry detection, a Z561/10-HC clean-up, HC TIRF dichroic, and 600/50-HC emission filter were used. All images were captured with Photometrics CoolSnap EZ monochrome CCD camera mounted to the microscope.

### Chemical treatments under fluorescence live-cell imaging

A stock solution of 40 mM hydrogen peroxide (H₂O₂) was prepared fresh, on the day of imaging, by diluting 9 M H₂O₂ (Sigma, Cat. #216763) in sterile water (vehicle). Cells were imaged before and after the addition of 5 µL of 40 mM H₂O₂ (i.e., a final concentration of 100 µM). Also, a stock solution of 500 mM oxidized glutathione (GSSG; Sigma, Cat. #G4376) or reduced glutathione (GSH; Sigma, Cat. #G6013) in 300 mM HEPES (vehicle) was prepared fresh, on the day of imaging, and its pH was adjusted to 7.4 with 1M NaOH dissolved in 300 mM HEPES. In the case of GSSG, cells were imaged before and after the addition of 40 µL of 500 mM GSSG (i.e., a final concentration of 10 mM). In the case of GSH, however, cells were treated with 40 µL of 500 mM GSH (i.e., a final concentration of 10 mM) or vehicle (HEPES; a final concentration of 6 mM) for 16 hr prior to imaging.

### Seahorse metabolic flux analysis

∼20,000 Hs578T cells were seeded into 6 wells of Seahorse XFp Cell Culture Miniplate (Agilent, Cat. #103025-100) in the RPMI 1640 growth medium without gentamicin while two blank wells were filled with the same growth medium without cells. The 8-well miniplate was then incubated at 37 °C, 5% CO_2_ and 95% humidity for 24 hr. Afterward, cells were gently washed with a modified BSS (mBSS) solution containing 1 mM HEPES (pH 7.4) and incubated at 37 °C in a CO_2_-free incubator for 1 hr to allow for the equilibration of temperature and pH. The cells were gently washed once more with prewarmed (37 °C) mBSS solution, inspected for adherence, and loaded into Seahorse XF HS Mini instrument (Agilent). Assays were then performed according to the manufacturer’s protocol. Briefly, after basal ECAR measurements for four cycles (Mix: 3 min, Wait: 0 min, Measure: 3 min), 100 µM H_2_O_2_ or water was injected for experimental and control groups, respectively. Then, a mixture of 0.5 µM rotenone (Sigma, Cat # 557368) and 0.5 µM antimycin A (Sigma, Cat # A8674) was applied to both groups, followed by 50 mM 2-deoxyglucose (Sigma, Cat # D6134). Not only ECAR but also oxygen consumption rates (OCR) signals were continuously measured throughout the assay, which was performed in triplicate in each of three independent experiments.

### Western blot analysis

Hs578T cells were cultured in standard 6-well plates and transfected under conditions similar to those used for imaging. 7-8 hr after the start of transfection, the transfected cells were selected for 16-20 hr by adding 5 µg/mL puromycin. Puromycin-treated cells were harvested and lysed in the RIPA buffer (50 mM Tris–base (pH 7.5), 150 mM NaCl, 1 mM sodium EDTA, 1% NP-40, 0.5% sodium deoxycholate, and 0.1% SDS) that contains protease inhibitors (Thermo Fisher, Cat# A32953) and phosphatase inhibitors (Thermo Fisher, Cat# A32957). The bicinchoninic acid assay (Thermo Fisher, Cat# 23227) was subsequently used to determine protein concentrations in cell lysates [72]. We then prepared 7.5% SDS-PAGE gels and loaded 20 μg and 50 μg of each sample per lane for GSR and CAT detection, respectively. After gel electrophoresis (100 V for 2 hr), the BioRad Semi-Dry Transfer System was subsequently used to transfer resolved proteins to a 0.45 µm PVDF membrane (Thermo Fisher, Cat# 88518). The membrane was soaked in 5% nonfat milk prepared in 1×PBST (1.8 mM KH_2_PO_4_, 10 mM Na_2_HPO_4_, 2.7 mM KCl, 137 mM NaCl, and 0.1% Tween 20), followed by three washes with 1×PBST. Then, a primary antibody was incubated with the membrane overnight at 4 °C for hGSR and hCAT while 1 hr at room temperature for β-actin. The primary antibodies used were rabbit anti-GSR (1:1000; Invitrogen, Cat# PA5-29945), rabbit anti-CAT (1:1000; Invitrogen, Cat# 702732), and mouse anti-β-actin (1:5000, Sigma, Cat# A5441). Subsequently, a secondary antibody that was conjugated with a near-IR dye was incubated with the membrane for 1 h at room temperature, which were either donkey anti-rabbit-IRDye® 680RD (1:10000, LICORbio, Cat# 926-68073) or goat anti-mouse-DyLight680 (1:10000, Cell Signaling, Cat# 5470). LI-COR Odyssey Sa Near-IR imager (LI-COR) was used to scan the membrane, and ImageStudio v6.0 (LI-COR) was used for quantitative analysis.

### Generation of Hs578T-ecCAT cells stably expressing mEGFP-ecCAT

Hs578T cells grown in the DMEM growth medium were seeded into a standard 6-well plate and cultured until they reached ∼70% confluence. Cells were then transfected with a plasmid expressing mEGFP-ecCAT and allowed to recover in the DMEM growth medium without antibiotics for 24 hr. Afterward, the cells were placed in the DMEM selection medium containing DMEM, 10% FBS, 500 µg/mL G418 (Corning, Cat. # 30-234-CI) and 50 µg/mL gentamicin. The cells were maintained in the selection medium for ∼3 weeks, with fresh medium replaced every 3-4 days. Each well was then examined under a fluorescence microscope to assess the expression of mEGFP-ecCAT. A well with the highest mEGFP fluorescent signals was then selected for further growth and passage into a T25 culture flask. After ∼24 hr, the stably transfected cells were maintained in the DMEM growth medium containing 250 µg/mL G418 with subculturing every 3-4 days for at least 2 weeks. The stably transfected cells (i.e., Hs578T-ecCAT) were then switched to the RPMI 1640 growth medium with the addition of 250 µg/mL G418. After another 2 weeks, the stably transfected Hs578T-ecCAT cells were deemed ready for experiments. Note that ∼70% of Hs578T-ecCAT cells were found to stably express mEGFP-ecCAT.

### Image Analysis

Captured fluorescence images were analyzed using ImageJ/FIJI software (National Institutes of Health) as reported previously [25]. Briefly, images were first scaled according to the microscope’s pixel size (0.12 µm/pixel). A duplicate of each scaled image was then subjected to the Robust Automated Threshold Selection (RATS) algorithm using default parameters (noise threshold = 25, λ factor = 3). The ‘Analyze Particles’ function was used to determine the number and area of fluorescent particles within a single cell and to create a corresponding mask, which was then used to eliminate overlapping particle counts. This process was repeated for all subsequent cell images.

### Statistical Analysis

All statistical analyses were performed using GraphPad Prism 9. A two-way ANOVA was used to assess the statistical significance of the treatment versus control groups. Tukey’s multiple comparisons test was subsequently performed to identify significant differences between specific treatment and control groups under each category. Student’s *t*-test was also performed when appropriate. Statistical significance was defined as *p* < 0.05 with a 95% confidence interval: * *p* < 0.05; ** *p* < 0.01; *** *p* < 0.001; and **** *p* < 0.0001; ns, not significant.

## Acknowledgements

We would like to thank Dr. Brian Polster (University of Maryland, Baltimore) for assisting with Seahorse metabolic flux analysis. This work was funded in part by the National Institutes of Health; R01GM134086 (M.K.), R01GM125981 (S.A.), and T32GM158458 (B.B.K.). The content is solely the responsibility of the authors and does not necessarily represent the official views of the National Institutes of Health.

## Contributions

A.S.: Conceptualization, Data curation, Formal analysis, Investigation, Validation, Visualization, Writing – original draft, Writing – review and editing.

B.B.K.: Investigation, Writing – review and editing.

M.K.: Data curation, Supervision, Funding acquisition, Writing – review and editing.

S.A.: Project administration, Conceptualization, Funding acquisition, Data curation, Formal analysis, Supervision, Visualization, Writing – review and editing.

All authors reviewed and edited the manuscript.

## Data Availability

All data are available within the manuscript and the supplementary materials.

