## Supplemental Figures for "Functional Contribution of Multienzyme Glucosome Condensates to Cellular Redox Homeostasis in Cancer Cells"

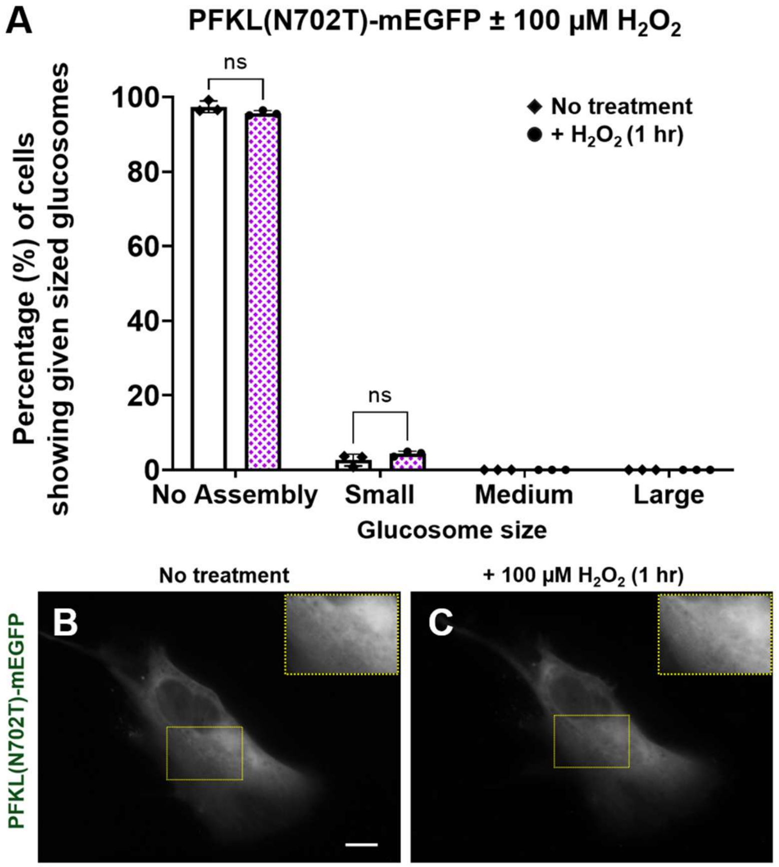


**Figure S1. $H_2O_2$ treatment of Hs578T cells expressing PFKL(N702T)-mEGFP.** *A*, the percentages (%) of cells showing given sized glucosomes were quantified before (blank bars, $N_{expt}$ = 3) and after (shaded bars, $N_{expt}$ = 3) the addition of $H_2O_2$ (100 µM, 1 hr). *B and C,* a representative Hs578T cell expressing PFKL(N702T)-mEGFP was imaged before (B) and after (C) the addition of $H_2O_2$ (100 µM). Insets indicate digitally magnified regions of interest. Scale bars, 10 µm. Error bars represent standard deviations (SD). Statistical analyses were performed using Tukey's multiple comparison tests for two-way ANOVA analysis. ns, not significant; $N_{expt}$, the number of independent experiments.

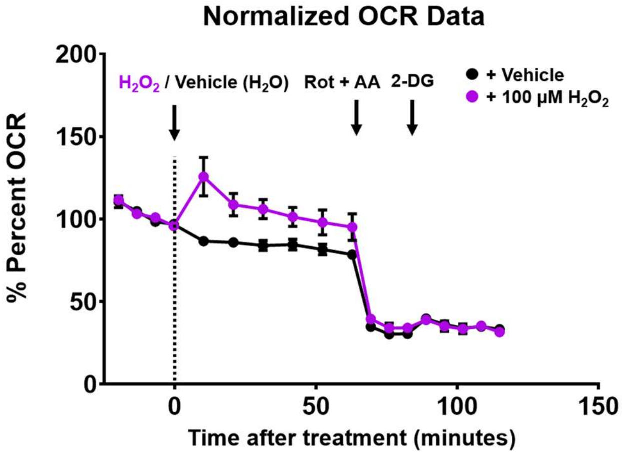


**Figure S2. Oxygen consumption rate (OCR) of Hs578T cells with $H_2O_2$.** Normalized % OCR of vehicle ($H_2O$, black line) and $H_2O_2$ (purple line) treated Hs578T cells shows a sharp decrease after the addition of rotenone (Rot) and antimycin A (AA). Black arrows indicate the timepoint for the addition of i) 100 µM $H_2O_2$ or vehicle ($H_2O$), ii) a mixture of 0.5 µM Rot and 0.5 µM AA, and iii) 50 mM 2-deoxyglucose (2-DG). Each data point represents average % OCR signals from three technical replicates of three independent trials with error bars representing standard deviations.

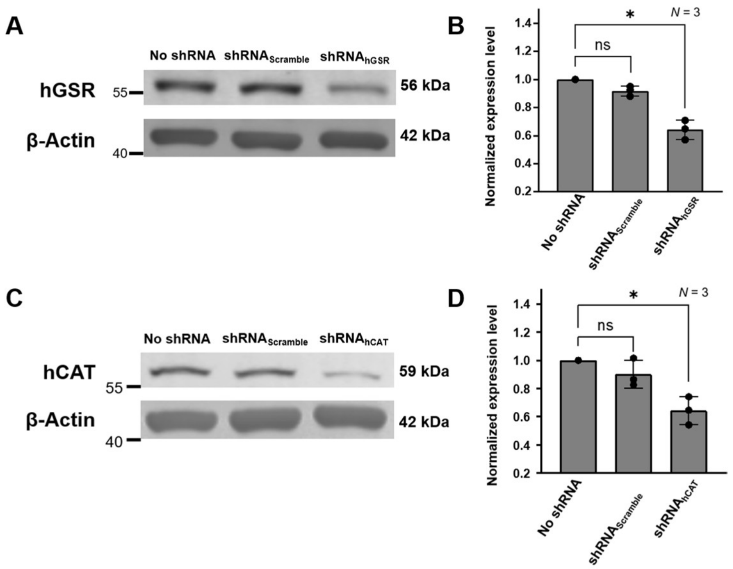


**Figure S3. Knockdown of human glutathione reductase (hGSR) and catalase (hCAT) in Hs578T cells.** Western blot analyses show the knockdown of hGSR (A and B) and hCAT (C and D) in Hs578T cells, respectively. Their expression levels were normalized based on β-actin. Error bars represent standard deviations (SDs) from three independent experiments. Statistical analyses were performed using Student's *t*-test. * $p < 0.05$; ns, not significant.

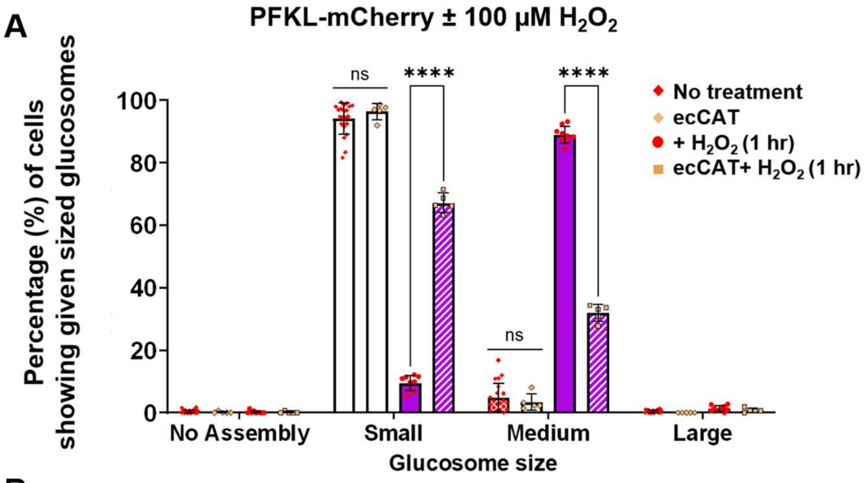


**B**

| Cell line | Conditions | Percentage (%) of cells displaying given sized glucosomes | | | |
|---|---|---|---|---|---|
| | | No Assembly | Small | Medium | Large |
| Hs578T | No treatment * | 0.4 ± 0.6 | 94.3 ± 4.7 | 4.9 ± 4.6 | 0.4 ± 0.4 |
| | + 100 μM $H_2O_2$* | 0.2 ± 0.5 | 9.4 ± 2.4 | 89.0 ± 2.7 | 1.4 ± 0.9 |
| Hs578T-ecCAT | No treatment | 0.2 ± 0.3 | 96.4 ± 2.6 | 3.4 ± 2.6 | ND |
| | + 100 μM $H_2O_2$ | 0.1 ± 0.3 | 67.2 ± 3.1 | 31.9 ± 2.7 | 0.8 ± 0.7 |

* Data from Figure 1F for direct comparison. ND = Not Detected

**Figure S4. Expression of *E. coli* catalase (ecCAT) in Hs578T cells reduces the effect of $H_2O_2$ on glucosome formation.** *A,* the percentages (%) of Hs578T cells showing given sized glucosomes with and without stably expressing ecCAT were imaged before (blank bars, $N_{expt}$ = 5) and after (shaded bars, $N_{expt}$ = 5) the addition of $H_2O_2$ (100 µM, 1 hr). Note that the data from **Figure 1F** of Hs578T cells expressing PFKL-mCherry are also included as 'No treatment' and '+ $H_2O_2$ (1 hr)' in the graph for direct comparison. Error bars represent standard deviations (SD). *B*, a table shows the average percentages (%) of Hs578T cells showing given sized glucosomes with and without stably expressing ecCAT, and with or without $H_2O_2$, along with their SDs (±). Statistical analyses were performed using Tukey's multiple comparison tests for two-way ANOVA analysis. **** $p < 0.0001$; ns, not significant; $N_{expt}$, the number of independent experiments.

A

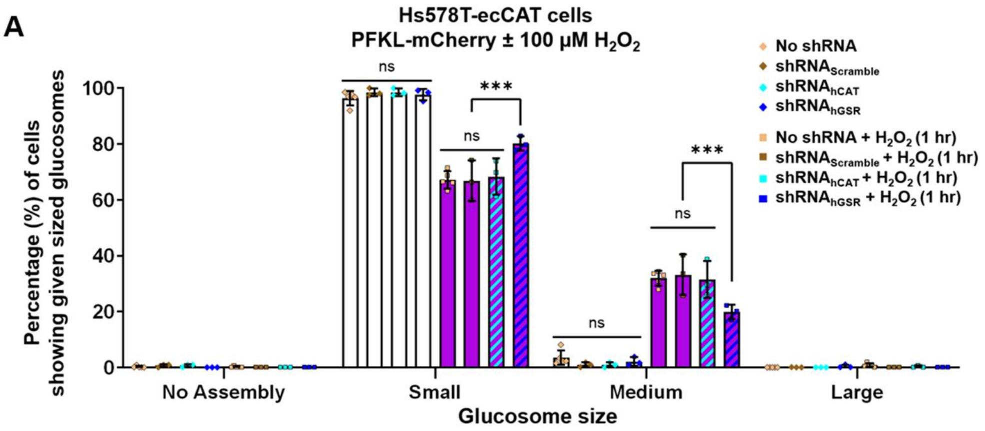


B

| Conditions | | Percentage (%) of cells displaying given sized glucosomes | | | |
|---|---|---|---|---|---|
| | | No Assembly | Small | Medium | Large |
| No Treatment | No shRNA* | 0.2 ± 0.3 | 96.4 ± 2.6 | 3.4 ± 2.6 | ND |
| | $shRNA_{Scramble}$ | ND | 99.2 ± 0.8 | 0.3 ± 0.5 | 0.5 ± 0.5 |
| | $shRNA_{hCAT}$ | 0.6 ± 0.5 | 98.5 ± 1.4 | 0.9 ± 0.9 | ND |
| | $shRNA_{hGSR}$ | ND | 97.7 ± 2.0 | 2.0 ± 1.5 | 0.3 ± 0.5 |
| + $H_2O_2$ (1 hr) | No shRNA* | 0.1 ± 0.3 | 67.2 ± 3.1 | 31.9 ± 2.7 | 0.8 ± 0.7 |
| | $shRNA_{Scramble}$ | ND | 66.8 ± 7.3 | 33.2 ± 7.3 | ND |
| | $shRNA_{hCAT}$ | ND | 68.3 ± 6.5 | 31.5 ± 6.6 | 0.2 ± 0.4 |
| | $shRNA_{hGSR}$ | ND | 80.2 ± 2.6 | 19.8 ± 2.6 | ND |

* Data from Figure S4 for direct comparison. ND = Not Detected

**Figure S5. Knockdown effect of human catalase (hCAT) and glutathione reductase (hGSR) on $H_2O_2$-induced glucosomes in Hs578T-ecCAT cells**. *A,* the percentages (%) of Hs578T cells showing given-sized glucosomes with $shRNA_{Scramble}$, $shRNA_{hCAT}$, or $shRNA_{hGSR}$ were quantified before (blank bars, $N_{expt}$ = 3, $N_{expt}$ = 3, and $N_{expt}$ = 3, respectively) and after (colored bars, $N_{expt}$ = 3, $N_{expt}$ = 3, and $N_{expt}$ = 3, respectively) the addition of $H_2O_2$ (100 µM, 1 hr). Note that the data from **Figure S4** of Hs578T-ecCAT cells expressing PFKL-mCherry (i.e., 'ecCAT' and 'ecCAT+ $H_2O_2$ (1 hr)') are used here as 'No shRNA' and 'No shRNA+ $H_2O_2$ (1 hr)' respectively for direct comparison. Error bars represent standard deviations (SD). *B*, a table shows the average percentages (%) of Hs578T cells showing given sized glucosomes in various conditions along with their SDs (±). Statistical analyses were performed using Tukey's multiple comparison tests for two-way ANOVA analysis. *** $p < 0.001$; ns, not significant; $N_{expt}$, the number of independent experiments.
